# Vitamin B5, a Coenzyme A precursor, rescues TANGO2 deficiency disease-associated defects in *Drosophila* and human cells

**DOI:** 10.1101/2022.11.04.514597

**Authors:** Paria Asadi, Miroslav P. Milev, Djenann Saint-Dic, Chiara Gamberi, Michael Sacher

**Affiliations:** Concordia University, Department of Biology, Montreal, Quebec, Canada, H4B1R6; Coastal Carolina University, Department of Biology, Conway, South Carolina, USA, 29526; McGill University, Department of Anatomy and Cell Biology, Montreal, Quebec, Canada, H3A0C7

**Keywords:** TANGO2, vitamin B5, membrane traffic, coenzyme A, metabolic crisis, neurodevelopment, *Drosophila*

## Abstract

Mutations in the <u>T</u>ransport <u>an</u>d <u>G</u>olgi <u>O</u>rganization 2 (*TANGO2*) gene are associated with intellectual deficit, neurodevelopmental delay and regression. Individuals can also present with an acute metabolic crisis that includes rhabdomyolysis, cardiomyopathy and cardiac arrhythmias, the latter of which are potentially lethal. While preventing metabolic crises has the potential to reduce mortality, no treatments currently exist for this condition. The function of TANGO2 remains unknown but is suspected to be involved in some aspect of lipid metabolism. Here, we describe a model of *TANGO2*-related disease in the fruit fly *Drosophila melanogaster* that recapitulates crucial disease traits. Pairing a new fly model with human cells, we examined the effects of vitamin B5, a Coenzyme A (CoA) precursor, on alleviating the cellular and organismal defects associated with *TANGO2* deficiency. We demonstrate that vitamin B5 specifically improves multiple defects associated with TANGO2 loss-of-function in *Drosophila* and rescues membrane trafficking defects in human cells. We also observed a partial rescue of one of the fly defects by vitamin B3, though to a lesser extent than vitamin B5. Our data suggest that a B complex supplement containing vitamin B5/pantothenate may have therapeutic benefits in individuals with TANGO2-deficiency disease. Possible mechanisms for the rescue are discussed including restoration of lipid homeostasis.

**SYNOPSIS:** Using a *Drosophila* fruit fly model that recapitulates many defective phenotypes associated with TANGO2 deficiency disease (TDD), we show that treatment with vitamin B5 rescues these defects and suggest a multivitamin or B complex vitamin containing vitamin B5 may prevent the potentially lethal metabolic crises associated with TDD.

## INTRODUCTION

Initially identified nearly two decades ago as a protein that affects membrane trafficking in insect cells,^1^ TANGO2 is conserved from the fruit fly *Drosophila melanogaster* to humans and was only recently linked to human health.^2, 3^ Affected individuals with TANGO2-deficiency disease (TDD) manifest a plethora of clinical features (summarized in Schymick et al ^4^) including neurodevelopmental delay, seizures, developmental regression and intellectual deficit. Acute metabolic crises, often triggered by fever or fasting, manifest as, hypoglycemia, lactic acidemia and elevated creatine phosphokinase and can lead to episodic rhabdomyolysis and potentially lethal cardiac arrhythmias.^5^ Little is known about the function and localization of the TANGO2 protein with studies suggesting mitochondrial, cytosolic or a mix of both localizations.^2, 3, 6, 7, 8^ Several reports have shown defective endomembrane trafficking and dilated endoplasmic reticulum (ER) in fibroblasts derived from TDD affected individuals. ^3, 7^ Other studies have demonstrated defective mitochondrial morphology and respiration ^3, 7, 9^ leading to the speculation that TANGO2 may affect some as yet unknown aspect of lipid metabolism One study reported abnormal palmitate oxidation in cells derived from two individuals harboring two different *TANGO2* alleles, suggesting a defect in fatty acid oxidation and/or oxidative phosphorylation.^2^ These various results have made it difficult to devise treatment options for affected individuals, particularly to avoid the metabolic crisis which is the sole contributor of TDD lethality.

We hypothesized that if TANGO2 participates in lipid metabolism, then increasing the levels of the fatty acid activator Coenzyme A (CoA) might ameliorate cellular and organismal defects associated with TANGO2 loss of function. Pantothenic acid (also known as vitamin B5) is the obligate precursor of CoA in an essential pathway conserved in both prokaryotes and eukaryotes. In humans, the main source of pantothenic acid is dietary and a conserved, canonical five-step process converts this vitamin into CoA^10^, though a possible parallel pathway has also been reported in *Drosophila*.^11^ Both humans and *Drosophila* utilize four enzymes in the canonical pathway - three monofunctional enzymes (PANK, PPCS and PPC-DC) that carry out the first three steps, and one bifunctional enzyme (COASY) catalysing the final two steps^10^, - further highlighting the conservation of this pathway. The generated CoA is used in a variety of cellular processes including the TCA cycle, lipid activation for fatty acid oxidation and synthesis as well as lipoylation of specific proteins. Some of these cellular processes are carried out by holo-acyl carrier proteins (holoACP), which themselves require the 4’-phosphopantetheine group from CoA for their production.^12^ To test our hypothesis, we subjected two models of TDD to vitamin B5 supplementation and assessed phenotypic rescue. We report that the *Drosophila TANGO2*^*G517*^ loss-of-function mutants displayed several defects that overlap with the clinical phenotypes of TDD-affected individuals and can be used as a miniature whole-animal model of *TANGO2* dysfunction/loss-of-function. We found that vitamin B5 supplementation rescued the defective phenotypes of the *TANGO2*^*G517*^ mutant up to near wild type levels. These effects are specific to vitamin B5 since vitamins C, B3 and B9 supplementation did not result in a similar level of rescue, though vitamin B3 did show a modest effect in one assay. Continuous supplementation from larval development to adults was not required for rescue but positive effects appeared time-dependent. Similarly, in cultured human fibroblasts, vitamin B5 rescued a *TANGO2*-dependent cellular trafficking defect. Our results suggest that supplementation with a multivitamin or a B complex vitamin containing vitamin B5, and perhaps vitamin B3, may benefit individuals affected by TDD.

## METHODS

### Drosophila melanogaster maintenance, growth and vitamin supplementation

*TANGO2*^*G517*^ flies were obtained from the Bloomington *Drosophila* Stock Center. Control *Oregon*^*R*^ (*Ore*^*R*^) and *TANGO2*^*G517*^ fruit flies were maintained at 20-23°C on a Bloomington formula commercial fly food (Genesee Scientific). Flies aged 0-2 days old were collected and aged 10 days to yield the 10-12 days old populations to assay. Flies were kept well-fed by transferring to fresh food every three days. Supplemental vitamin B5 (Sigma, P5155) (2-4 mM as indicated) ^11^, vitamin B3 (Sigma, N0636) (0.5 mM) ^13^, vitamin B9 (Sigma, F7876) (10 µM) ^14^ or vitamin C (ThermoFisher, A61-100) (7 mM) ^15^ were dissolved in water and added to the warm liquid food and allowed to solidify at room temperature. Unused food was stored at room temperature for no longer than 10 days before use.

### Real time quantitative PCR (RT-qPCR)

Total RNA was collected from 50-200 fly heads using TRIzol™ reagent (ThermoFisher) as per the manufacturers protocol. Extracted RNA was converted to cDNA using the First Strand cDNA Synthesis kit (Origene) and then subjected to real time quantitative PCR using SYBR Green Master mix (BioRad) and the following oligonucleotides: TANGO2-forward 5’-GGCTGCTCAAAGATTGCACC-3’; TANGO2-reverse 5’-TTGCCAAATCCGTAGCACTCG-3’; Rpl32-forward 5’-AGCATACAGGCCCAAGATCG-3’; Rpl32-reverse 5’-TGTTGTCGATACCCTTGGGC-3’. The experiment was performed 3 times, with each cDNA sample amplified in triplicate. The TANGO2 expression was normalized to that of Rpl32.

### Drosophila assays

All assays were performed on larvae and on both male and female adults (10-12 days old) of the *Ore*^*R*^ and *TANGO*^*G517*^ genotypes. The fly *TANGO2* gene is on the X chromosome, and we noticed that there were much fewer males in the *TANGO2*^*G517*^ colony. The reason for this remains unclear. Recovering higher numbers of female flies, assays were first performed with females and are shown in the main results section Assays were then repeated with males and are shown in the supplemental information section. All assays were repeated a minimum of 3 times. Adults were anaesthetized with CO_2_ to separate males from females and allowed to reacclimate for 45 minutes prior to any assay. The total number of flies assayed from all biological replicates (N) is specified for each assay below. The N values for the specific experiments shown in each figure are indicated in the figure legends. The largest values represent controls which were repeated multiple times while testing various vitamin conditions. In each figure, one representative experiment is shown except for figures 1D, 2B, C and D (and supplemental figures 1D, 2B and C), for which cumulative results are displayed.

**Figure 1.**
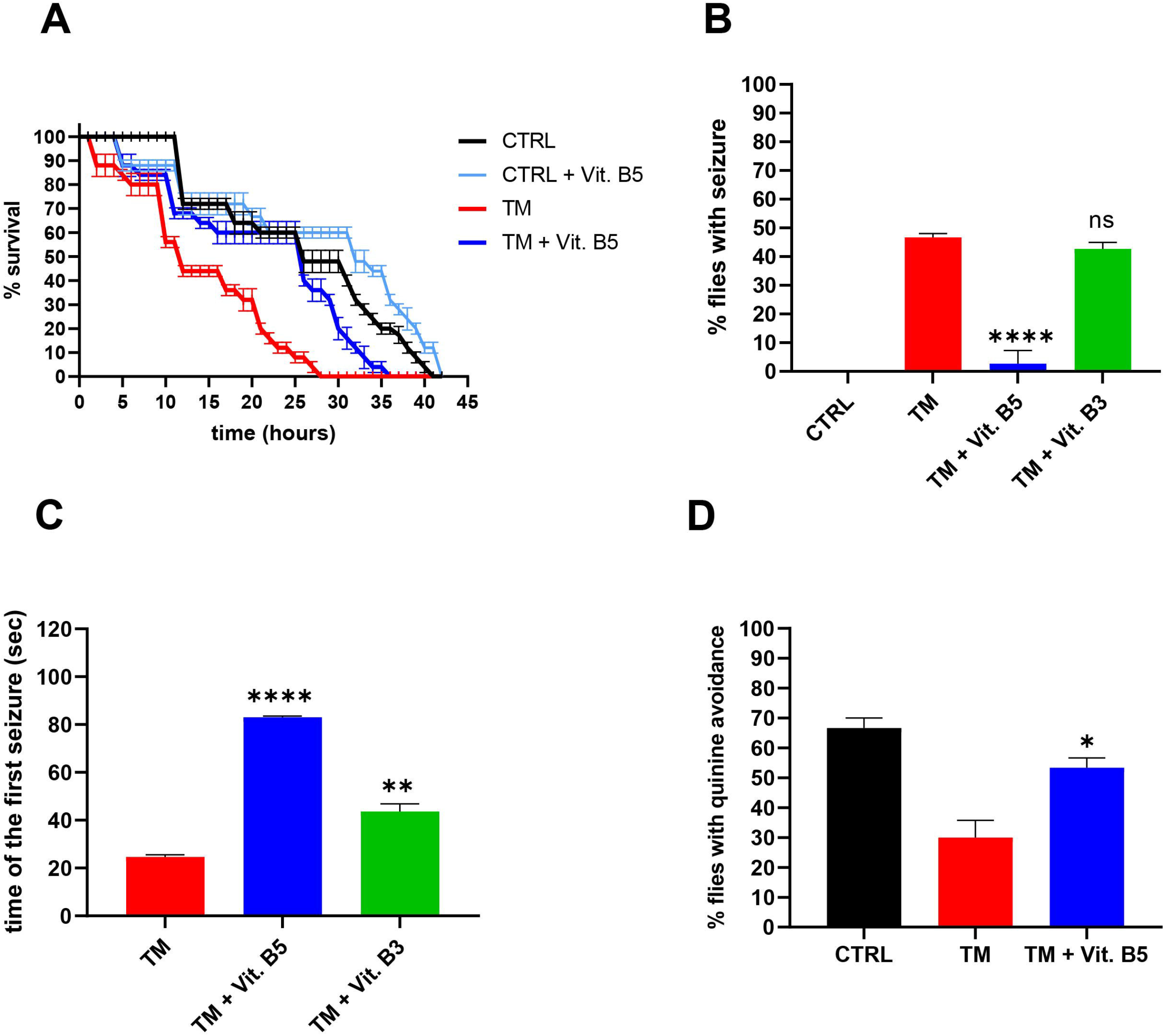
The *Drosophila TANGO2*^*G517*^ mutants display behavioral defects that are rescued by vitamin B5 supplementation. (A) Female fly populations were placed into empty plastic vials and survival was examined every hour until all flies were dead. (B), (C) Female fly populations were subjected to elevated temperatures for 120 seconds as detailed in the Methods section. The percentage of flies experiencing a seizure (B) and the time to the first seizure (C) are shown. (D) Female fly populations were subjected to a learning assay as described in the Methods section. The percentage of flies that learned to avoid the quinine-laced light side of the setup were recorded. Where indicated, supplements of either vitamin B5or B3 were included in the food for at least one generation prior to the assays. The concentrations of vitamin B5 used were 4 mM in (A) and 2 mM in (B), (C) and (D). N values for each experiment shown were 75 in (A) and (B), and (C), and 30-40 in (D). The asterisks for statistical analyses indicate p≤0.05 (*), p≤0.01 (**) and p≤0.0001 (****). CTRL represents *Ore*^*R*^ flies and TM represents *TANGO2*^*G517*^ flies. Results obtained with male flies are shown in Figure S1.

#### Starvation assay

Flies of age 10-12 days old were placed into empty plastic vials and starved for food and fluids. Survival was monitored hourly in 3 assay replicates. The total N values ranged from 750-3000.

#### Larvae motility assay

Third instar larvae were placed at the center of a 10 cm petri dish containing 2% agar in water. Larvae were observed over 9 minutes and the total crawling distance moved was recorded. The direction of movement was assessed by comparing the total distance moved from the initial position of larva on the dish (d) to the greatest displacement in any direction from the initial position (ΔX_max_). The total N values ranged from 15-60.

#### Climbing (geotaxis) assay

The protocol of Tapia et al ^16^ was modified as follows. The target line was placed 5 cm from the bottom of the tube and flies were given 10 seconds to reach this target line. The total N values ranged from 225-2500.

#### Open field assay

The protocol of Tapia et al ^16^ was followed except that the distance moved was recorded manually. Using a 10 cm diameter dish, the field was set as a 5 cm diameter from the center of the dish. The total N values ranged from 15-20.

#### Learning assay

Flies were placed on 1% agarose in water for 16 hours before acclimating for 10 minutes in the dark. The protocol of Tapia et al ^16^ was modified as follows. A disk of filter paper wetted in 0.1M quinine (a bitter substance that repulses flies) was placed in the lighted end of the setup, and the flies were allowed 30 seconds to venture into the lighted end of the setup. The total N values ranged from 40-50.

#### Seizure induction

Flies were acclimated in an empty plastic vial for 45 minutes prior to the assay. The vials were then immersed in a beaker containing water at either 42°C (males) or 42.3°C (females) for 120 seconds and the time to the first seizure was noted. Flies counted to be experiencing a seizure were those who fell to the bottom of the tube and underwent random spasms. The total N values ranged from 225-300.

Statistical analyses were performed using GraphPad Prism as one-way ANOVA with a post-hoc Tukey HSD test. Data are shown with S.E.M. error bars.

### Membrane trafficking assay

The retention using selective hooks (RUSH) assay was performed as previously described.^7^ Human fibroblasts grown in DMEM with 10% fetal bovine serum were treated with 2 mM vitamin B5, 0.5 mM vitamin B3 or 0.5 mM vitamin C for four days prior to performing the assay. The assay was performed a minimum of three times for each condition.

## RESULTS

### TANGO2^G517^ mutant flies display multiple behavioral defects

The human (276 amino acids) and *Drosophila* (283 amino acids) TANGO2 proteins have 30% identity and 50% similarity over their entire length. The predicted structures based on alphaFold ^17^ are also similar, with the pairwise alignment giving a root mean square distance value of 1.83 and a template modeling score of 0.9 over 96% coverage. We therefore evaluated *Drosophila* as a model organism to study TDD. The *TANGO2*^*G517*^ allele has a P-element construct inserted at the 5’ end of the *TANGO2* gene. Although these flies are viable, they are delayed in development relative to the *Ore*^*R*^ wild type control. *TANGO2* expression was measured by quantitative real-time PCR from RNA extracted from heads which showed that the *TANGO2*^*G517*^ flies had a residual 3.13%±1.23 expression compared to that of wild type *Ore*^*R*^ control flies of equal age. Thus, the *TANGO2*^*G517*^ mutant flies can serve as a good model to examine the effects of TANGO2 loss-of-function.

We initially performed a series of assays that replicate triggers of metabolic crises in TDD individuals and assays that measure some of the common non-neuromuscular defects. Since starvation triggers a metabolic crisis in TDD individuals^18^, we first examined the viability of 10-12 days old adult flies upon starvation. Both *Ore*^*R*^ and *TANGO2*^*G517*^ flies were starved for both food and moisture by placing them in empty vials. The *TANGO2*^*G517*^ flies died faster than *Ore*^*R*^ wild type flies with 50% viability reached at ~12 hours for the former and ~25 hours for the latter (Figures 1A and S1A).

A common feature of individuals suffering from TDD is seizures. We therefore assessed the heat-induced seizure rate of *TANGO2*^*G517*^ flies compared to *Ore*^*R*^ wild type. While the *Ore*^*R*^ wild types did not experience seizures due to the elevated temperatures, ~40% of *TANGO2*^*G517*^ flies did (Figures 1B and S1B), with the first seizure occurring at ~30 seconds (Figures 1C and S1C). Female flies required slightly higher temperatures (42.3°C) in order to induce seizures in the *TANGO2*^*G517*^ flies compared to males (42°C).

Since intellectual deficit is a common feature of TDD, we assessed the fly learning ability.^16, 19^ In this assay, flies are kept in the dark and then allowed to move toward a bright area of the housing vial following their natural attraction to light. The bright end of the vial was, however, laced with quinine, a bitter substance repulsive to flies. The flies were then reacclimated for 10 seconds on the dark side of the vial and allowed to venture into the light side six more times before finally testing their ability to avoid the quinine-laced side over a 10-second period. While ~65% of the *Ore*^*R*^ flies learned to avoid the repulsive lighted side, only about 30% of the female *TANGO2*^*G517*^ flies did (Figures 1D and S1D). We also noted that *Ore*^*R*^ flies already showed signs of learning during early phases of the training sessions (trials 1-5) prior to the actual test (trial 7) whereas it took longer for the *TANGO2*^*G517*^ flies to achieve avoidance (Figures S1E and F).

We next performed a series of assays to test possible motor defects. First, we examined geotaxis of the 10–12-days old adults by gently jolting them to the bottom of an empty vial and assessing the rate at which they climb 5 cm to a target line. As above, the *TANGO2*^*G517*^ flies performed worse than control, with an ~80% reduction of climbing compared to *Ore*^*R*^ (Figures 2A and S2A).

**Figure 2.**
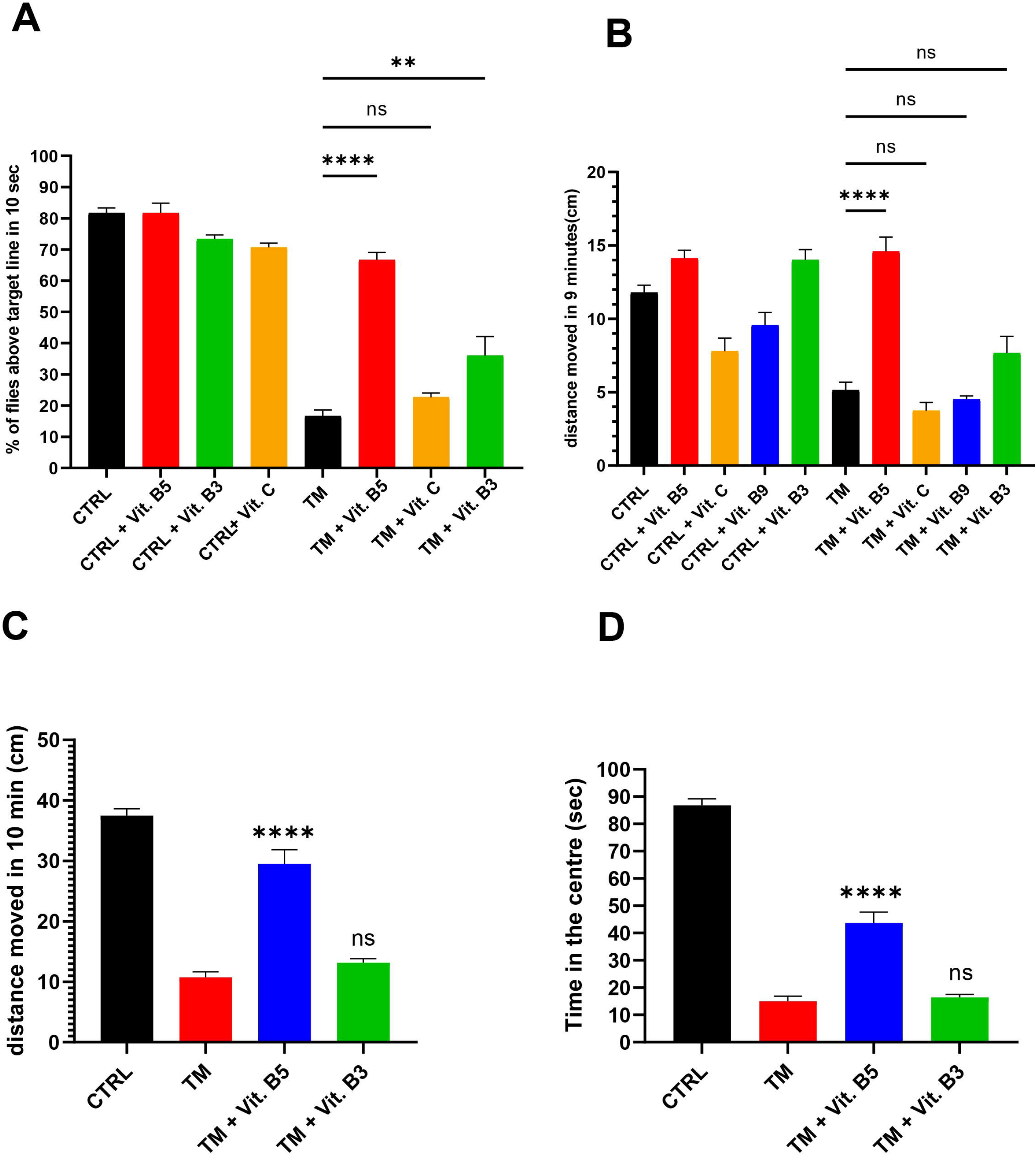
Motor defects associated with the *TANGO2*^*G517*^ mutants are rescued by vitamin B5 supplementation. (A) Female flies were placed into an empty vial, gently knocked to the bottom of the vial and were examined as they climbed. The percentages of flies that reached a line 5 cm from the bottom of the vial within 10 seconds are shown. (B) Third instar larvae were observed as they crawled over 9 minutes on a dish containing 2% agar. The total distance moved is shown. (C) Adult females were placed in the center of a 10 cm diameter petri dish and the distance moved over 10 minutes is displayed, as well as the time spent in the central area of the dish (D). Vitamins B5, B3, B9 or C as indicated were included in the food for at least one generation prior to the assays. The concentrations of vitamin B5 used were 4 mM in (A), (C) and (D), and 2 mM in (B). N values for each experiment shown were 50-75 in (A), 15-35 in (B) and 15 in (C) and (D). The asterisks for statistical analyses indicate p≤0.01 (**) and p≤0.0001 (****). CTRL represents *Ore*^*R*^ flies and TM represents *TANGO2*^*G517*^ flies. Results for male flies are shown in Figure S2.

We then examined third instar larvae for their ability to move on agar medium over 9 minutes. As was seen with the adults, the *TANGO2*^*G517*^ larvae displayed reduced movement by greater than 50% compared to wild type (Figure 2B). In addition, while the wild type larvae tended to move in a straight line, the *TANGO2*^*G517*^ larvae moved in a more circular, and seemingly uncoordinated manner. This may be reminiscent of a form of ataxia often seen in individuals with TDD.^4^ In order to quantify this phenotype, we reasoned that larvae that move in a straight line will have a “ total distance moved/maximum magnitude of displacement from the initial point” (d/Δx_max_) ratio of ~1. Mutant larvae with circular motions would remain closer to the initial point despite the total distance moved. Hence, d/Δx_max_ would be >1. Indeed, we found that *Ore*^*R*^ larvae had a d/Δx_max_ ratio of 1.036±0.006 while *TANGO2*^*G517*^ larvae showed a significantly higher ratio of 1.378±0.076 (p<0.0001).

Open-field assays are used to assess anxiety in flies and other model organisms.^16, 20^ We evaluated 10-12 days old flies for their ability to walk on a plastic culture dish and monitored their location either within a 5 cm diameter area from the centre of the dish or peripherally to it. The *TANGO2*^*G517*^ flies walked on average ~75% shorter distances over the given time period compared to *Ore*^*R*^ wild type (Figures 2C and S2B). Interestingly, the *TANGO2*^*G517*^ flies showed a propensity to move toward the edge of the dish and linger, spending ~85% less time in the center of the dish than the *Ore*^*R*^ wild type (Figures 2D and S2C). Therefore, *TANGO2*^*G517*^ may exhibit anxious behaviour.

### Vitamin B5 reverses all of the behavioral defects in TANGO2^G517^ flies

Having observed a number of defects reminiscent of TDD in *TANGO2*^*G517*^ mutants, and since TANGO2 defects are thought to impair lipid metabolism, we sought to determine possible benefits of supporting lipid metabolism via vitamin B5 supplementation since this molecule is a precursor to the lipid activator CoA. We first titrated vitamin B5 levels, identifying 8 mM as the highest concentration that did not appear to negatively impact fly cultures (not shown). This amount is similar to that used in another study.^11^ Using the relatively fast geotaxis assay (shown in Figures 2A and S2A), we found that 2-4 mM vitamin B5 gave more robust results and settled on this range for all of the rescue experiments described below. In all cases, we found that vitamin B5 supplementation improved assay performance of the *TANGO2*^*G517*^ flies. Compared to untreated siblings, the female 10-12 days old *TANGO2*^*G517*^ flies fed vitamin B5 were able to survive longer upon starvation (50% survival at ~25 hours compared to 12 hours for vehicle-treated siblings flies) (Figure 1A), had significantreduction (~95% reduction) of heat-induced seizures and ~3.5 fold longer times before experiencing the first seizure (Figures 1B,C), performed nearly 2 fold better in the learning assay (Figures 1D and S1F), had ~3.5 fold stronger geotactic response (Figure 2A), walked ~3 fold further (Figure 2C), and stayed in the center of the culture dish ~3 fold longer (Figure 2D). Similar effects of vitamin B5 were seen in the male flies (Figures S1A, B, C, D and E, Figures S2A, B and C). The third instar larvae also showed increased movement after vitamin B5 supplementation compared to untreated larvae (Figure 2B), and regained the ability to move in a straight line rather than the uncoordinated circular movement typical of untreated siblings (d/Δx_max_ ratio of 1.120±0.022 for vitamin B5-treated larvae compared to 1.378±0.076 for untreated larvae (p<0.001)).

To determine if these effects were specific to vitamin B5, we tested other vitamins including two other B vitamins (B3 and B9) as well as vitamin C. While vitamins B9 and C did not improve larval crawling, seizures, and open field tests respectively (Figures 1B,C and 2B,C,D), vitamin B3 displayed a small but significant improvement of motor activity in the geotaxis assay (Figure 2A).

### Vitamin B5 shows a time-dependent rescue of behavioral defects

In the previous assays, flies were receiving vitamin B5 supplementation throughout development from the egg stage until 10-12 days old when adults were assayed. We next asked if adult *TANGO2*^*G517*^ flies grown in normal conditions could benefit from vitamin B5 supplementation over shorter periods of time. To address this, we used the geotaxis assay, the starvation assay and the heat-induced seizure assay that recapitulate critical features of TDD. The *TANGO2*^*G517*^ flies tested were fed either regular food (untreated) or food supplemented with vitamin B5 since the egg stage until the 10-12 days old age when adults were assayed, or grown in regular food and fed vitamin B5-supplemented food 1, 2 or 3 days prior to the assay. The *TANGO2*^*G517*^ flies showed a time-dependent increase in geotaxis (Figures 3A and S3A) and survival upon starvation (Figure 3B), and reduced incidence of heat-induced seizures upon vitamin B5 supplementation (Figures 3C and S3B). In each case, growth on vitamin B5 for as little as one day significantly rescued the *TANGO2*^*G517*^ defects, suggesting that although prolonged administration may prove most effective, vitamin B5 may exert benefits over a relatively short time frame.

**Figure 3.**
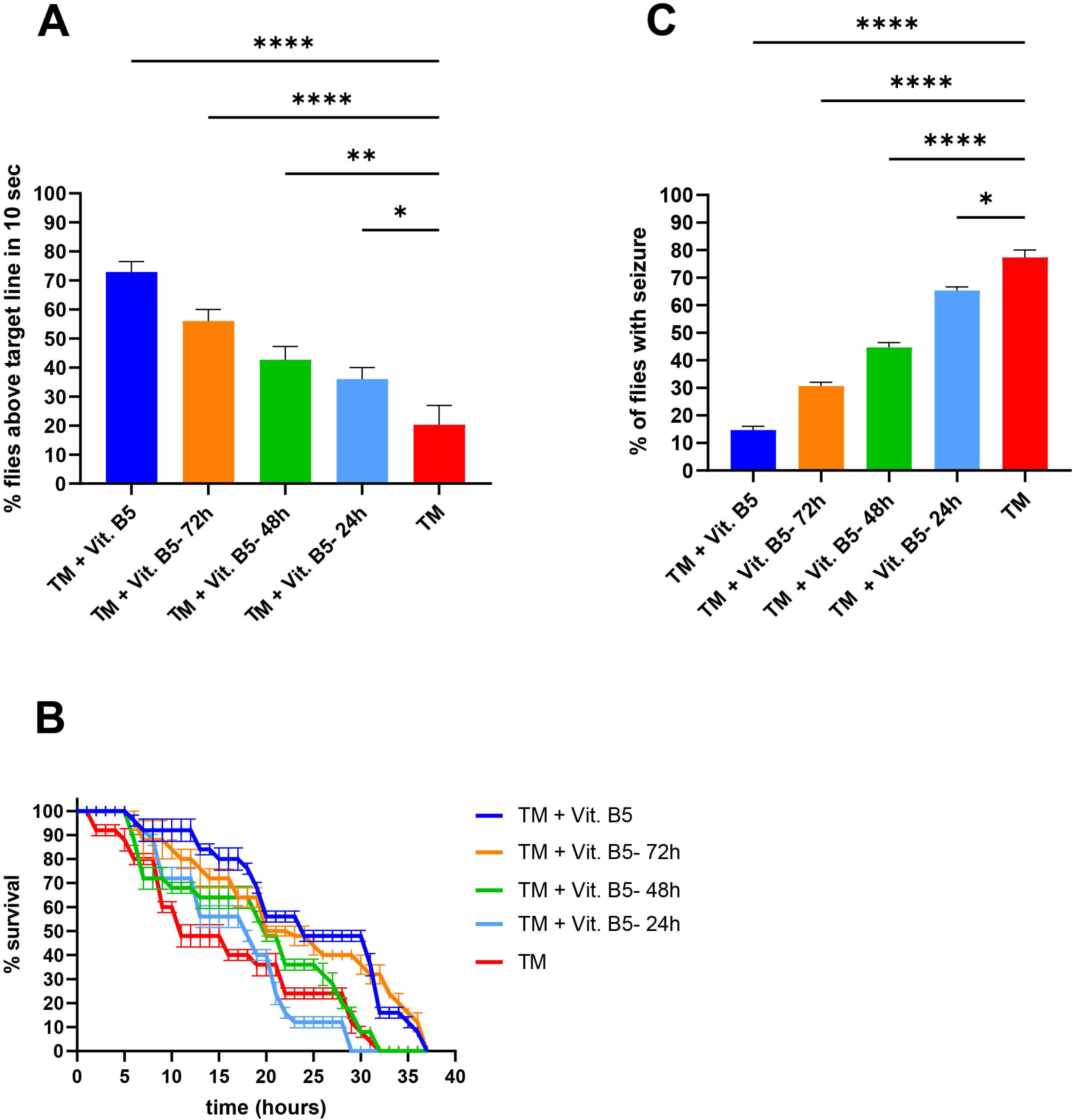
Vitamin B5 supplementation shows a time-dependent rescue of phenotypes in the *TANGO2*^*G517*^ mutants. *TANGO2*^*G517*^ female flies were grown either on food without vitamin B5, or grown for at least one generation either on regular food or on vitamin B5-supplemented food. A portion of the untreated *TANGO2*^*G517*^ adult flies were placed on vitamin B5-containing food either 1, 2 or 3 days prior to the assays. Flies were then assayed for their climbing ability (A), survival during starvation (B) and resistance to heat-induced seizures (C). The concentrations of vitamin B5 used were 4 mM in (A) and (C), and 2 mM in (B). N values for each experiment shown were 50-75 in (A), 75 in (B) and 75 in (C). The asterisks for statistical analyses indicate p≤0.05 (*), p≤0.01 (**), p≤0.001 (***) and p≤0.0001 (****). TM represents *TANGO2*^*G517*^ flies. Results obtained with male flies are shown in Figure S3.

### Vitamin B5 rescues a membrane trafficking defect in human fibroblasts devoid of TANGO2

Given the wide range of rescuing effects of vitamin B5 in the TDD *Drosophila* model, we asked if similar benefits would be found in mammals. We examined fibroblasts from a compound heterozygous individual with the common *TANGO2* exon 3-9 deletion and *TANGO2* exon 6 deletion, effectively rendering the cells a TANGO2 knockout ^2, 7^, and compared them to control fibroblasts. We first empirically determined that 2 mM vitamin B5 and 0.5 mM vitamin B3 supplementation in the growth medium for up to 4 days had no negative effects on the growth or morphology of the fibroblasts (not shown). Using an established assay for endoplasmic reticulum (ER) to Golgi transport, and as previously reported ^7^, we found the fibroblasts with the *TANGO2* alleles to have a greatly reduced rate of transport of the cargo protein from the ER to the Golgi (Figure 4A). Following vitamin B5 supplementation, the rate of transport was greatly increased to near control levels. The level of rescue was similar to that seen when the cells were rescued with wild type TANGO2 ^7^, suggesting that vitamin B5 could functionally replace TANGO2 in this assay. In these cells one allele (exon 3-9 del) is effectively a TANGO2 knockout, while the other allele (exon 6 del) could theoretically produce a truncated protein of ~15kD. Thus, it was formally possible that vitamin B5 could increase expression of the ~15kD truncation which might explain the rescue observed. However, as shown in Figure 4B, vitamin B5 treatment of the TANGO2 deficient fibroblasts did not result in the appearance of truncated TANGO2 protein. This suggests that the mechanism of action of vitamin B5, at least with respect to TANGO2, does not seem to involve an induction in protein expression. Corroborating the conclusion that vitamin B5 specifically rescues the trafficking defect of TANGO2 deficiency, neither vitamin B3 nor vitamin C supplementation showed a similar augmented rate of trafficking (Figure 4A). Collectively, our study shows that vitamin B5 supplementation can rescue a wide range of defects associated with *TANGO2* loss-of-function in TANGO2-deficient cells and whole organisms from *Drosophila* to humans.

**Figure 4.**
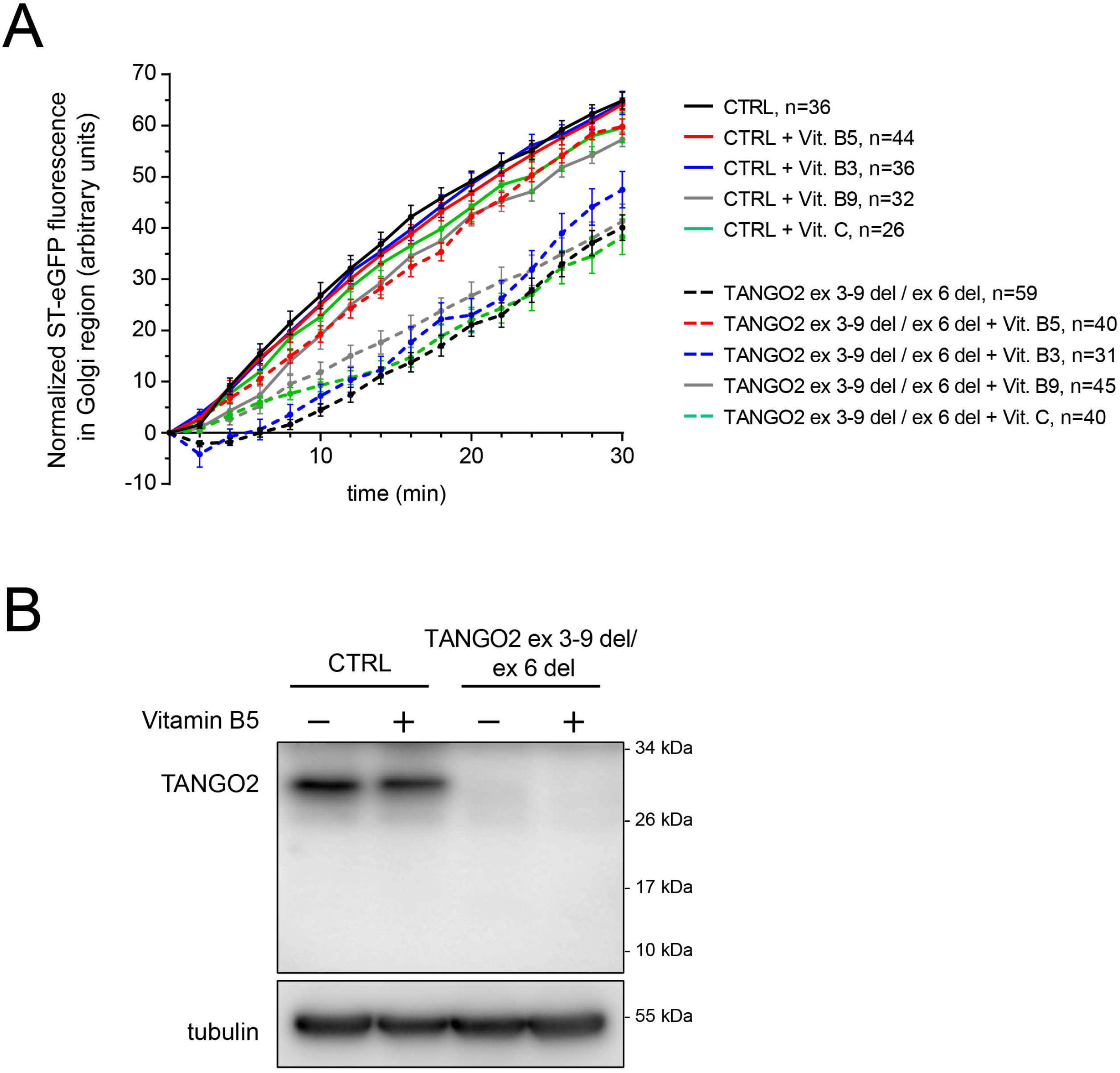
Vitamin B5 rescues an endoplasmic reticulum-to-Golgi trafficking defect in *TANGO2*-deficient human fibroblasts. (A) Wild type (CTRL) fibroblasts or fibroblasts harboring the compound heterozygous exon3-9 del/exon 6 del allele (*TANGO2*-deficient cells) were transfected with a plasmid expressing sialyltransferase-GFP-streptavidin binding protein which is retained in the ER by association with an ER-localized hook composed of streptavidin. After release of the GFP-tagged protein by biotin, fluorescence was monitored in the Golgi over time and quantified. Solid lines represent CTRL cells and dashed lines represent the *TANGO2* mutant cells. Black colouring represents untreated cells, red colouring represents vitamin B5-treated cells, blue colouring represents vitamin B3-treated cells and green colouring represents vitamin C-treated cells. Note that the only dashed line that clusters with the solid lines is from the vitamin B5-treated cells. (B) Lysates from CTRL and TANGO2-deficient cells either untreated or treated with vitamin B5 for 4 days were probed for TANGO2 and tubulin as a loading control. Note that the protein product of the exon 6 del allele would be expected to appear at ~15kD if it was produced and stable.

## DISCUSSION

TDD is a multisystem organ disease that includes neuromuscular, developmental, and intellectual delays. Affected patients are susceptible to clinical deterioration when exposed to metabolic stressors such as fever or fasting. These episodes of metabolic crisis lead to worsening ataxia, muscle weakness, and cardiac arrhythmias as well as cardiomyopathy and are the leading cause of mortality in this disease. Here, we find that *Drosophila TANGO2*^*G517*^ mutants exhibit greatly reduced *TANGO2* expression and display defects that recapitulate many of these clinical features. Thus, these mutants may be used to model TDD. Interestingly, all of the phenotypic defects seen in the *TANGO2*^*G517*^ flies are significantly reversed, some to near wild type levels, by vitamin B5 supplementation. It is noteworthy that a motor function was also rescued, albeit to a much lower, but still significant, degree by vitamin B3 that was shown to have broad positive effects in several unrelated conditions.^21^

To date, there are no treatment options for individuals suffering from TDD. A recent study suggests that folic acid (vitamin B9) can restore normal rhythm in a patient-derived cardiomyocyte model system.^22^ Together with our present study this suggests that several B complex vitamins may have beneficial effects on various clinical aspects of this disease. Consistent with this notion, a natural history study of over 70 individuals suffering from TDD found that metabolic crises were not seen in individuals taking a multivitamin or a B complex supplement.^23^ Multivitamins typically contain B vitamins including B5. Thus, our work and the aforementioned studies strongly suggest that B complex supplementation with at least vitamins B5 and B9, and possibly B3, may offer substantial therapeutic benefits for those affected by TDD.

The mechanism whereby these B vitamins rescue the TANGO2-dependent defects is currently unknown and awaits a thorough understanding of the molecular function(s) of the TANGO2 protein. Based on the trafficking assay in human fibroblasts (Figure 4A) the most likely scenario is that production of a metabolite, and not increased production of the TANGO2 protein, rescues the critical ER-to-Golgi function. In light of vitamin B5 being a precursor to CoA, it is tempting to speculate that increased levels of an unknown CoA metabolite(s) lead to a downstream phenotypic rescue in the two model systems.

The five-step biosynthesis pathway converting vitamin B5 to CoA has been linked to human neurodegenerative diseases with brain iron accumulation (NBIA). Variants in *PANK2*, mediating the first step in the pathway, and *COASY*, regulating the last two steps, are linked to pantothenate kinase associated neurodegeneration (PKAN or NBIA1; OMIM 234200) and COASY protein-associated neurodegeneration (CoPAN or NBIA6; OMIM 618266). Both conditions are characterized by impaired *de novo* synthesis of CoA. While individuals with PKAN, CoPAN and many with TDD exhibit cognitive impairment and movement disorders, TDD-affected individuals do not demonstrate the pathognomonic finding of brain iron accumulation observed in PKAN and CoPAN individuals.

Based on these differences, we can speculate on several possible cellular processes downstream of CoA that may be affected in TDD individuals. Acyl-CoA acts as a source of fatty acid for β-oxidation in mitochondrial energy production as well as a substrate for protein acylation. The TANGO2 protein was originally thought to function in membrane trafficking and it is noteworthy that this process is regulated by protein acylation^24^ suggesting this as a possible affected process in TDD. Another possibility is that TANGO2 acts downstream of holoACP, formed from apoACP by the transfer of the 4’-phosphopantetheine from CoA. holoACP is involved in fatty acid synthesis, defects in which could affect lipid homeostasis and the function of multiple membranes, a process that would respond to the dietary changes that trigger metabolic crises. Consistent with this notion are studies reporting minimal^18^, modest^2, 8^ and strong^25^ changes in acyl-carnitines in TDD-affected individuals. Further supporting this notion is a recent study by Malhotra and colleagues who demonstrated profound changes in the cellular lipid profile in a TANGO2 knockdown model which were further exaggerated upon starvation (Lujan et al, BioRxiv), consistent with preliminary lipidomic data from our group on fibroblasts devoid of TANGO2. holoACP is also involved in lipoylation of four protein complexes involved in mitochondrial and amino acid metabolism^26^, possibly explaining the reported oxidative phosphorylation defects and triggered metabolic crises. Alternatively, holoACP is involved in the synthesis of iron-sulfur clusters.^27^ In this respect it is noteworthy that a recent study suggests TANGO2 may act as a heme chaperone, moving this iron-containing molecule out of the mitochondria.^28^ If so, it remains to be determined how a TDD trigger relates to heme transport and a metabolic crisis, and why brain iron accumulation is not seen in TDD affected individuals.

Vitamin B3 is used in a salvage pathway to generate nicotinamide adenine dinucleotide (NAD). The latter molecule is a cofactor required in a number of cellular processes, perhaps best known as an electron acceptor/donor in energy production in the mitochondria. Given that several studies have suggested that a portion of the TANGO2 protein associates with mitochondria in humans and mice^7, 9, 29^, and a mitochondrial defect may be a secondary consequence of TANGO2 deficiency, the modest benefit of vitamin B3 in the climbing assay may reflect a non-specific overall increase in energy production or other factors, such as mitigation of oxidative stress. Consistent with this notion, vitamin B3 has been shown to be beneficial in a number of different, unrelated diseases.^21^

A precise definition of all TANGO2 cellular roles in the functional context of different tissues and organs will be needed to understand TDD and design targeted therapeutics. However, studies in several models support the conclusion that vitamins B5, B9 and perhaps B3 can alleviate and/or remove dangerous TDD manifestations even with a short course of administration. Considering the extensive knowledge about vitamin B supplementation and the minimal toxicity, these studies may improve the current care of individuals affected by TDD.

## Author contributions

PA performed and analyzed all of the *Drosophila* assays. MPM performed and analyzed the human fibroblast data. DSD performed qPCR and analyzed the data. CG conceived and guided the *Drosophila* work and analyzed the data. MS conceived the study, guided the work, analyzed the data and wrote the manuscript. All authors edited the manuscript and approved the final version.

## Ethics statement

Work on human fibroblasts has been approved by the Research Ethics committee of Concordia University.

## Data availability statement

he data supporting this study are available from the corresponding author upon reasonable request.

## Acknowledgements

We are grateful to members of the Sacher and Gamberi laboratories for critical review of this work, as well as to Drs. S. Mackenzie, L. Zhang, C. Miyake, S. Lalani and L. Ghaloul-Gonzalez for many helpful discussions. Stocks obtained from the Bloomington Drosophila Stock Center (NIH P40OD018537) were used in this study. Imaging was performed at the Centre for Microscopy and Cellular Imaging at Concordia University.

## FIGURE LEGENDS

**Figure S1.**
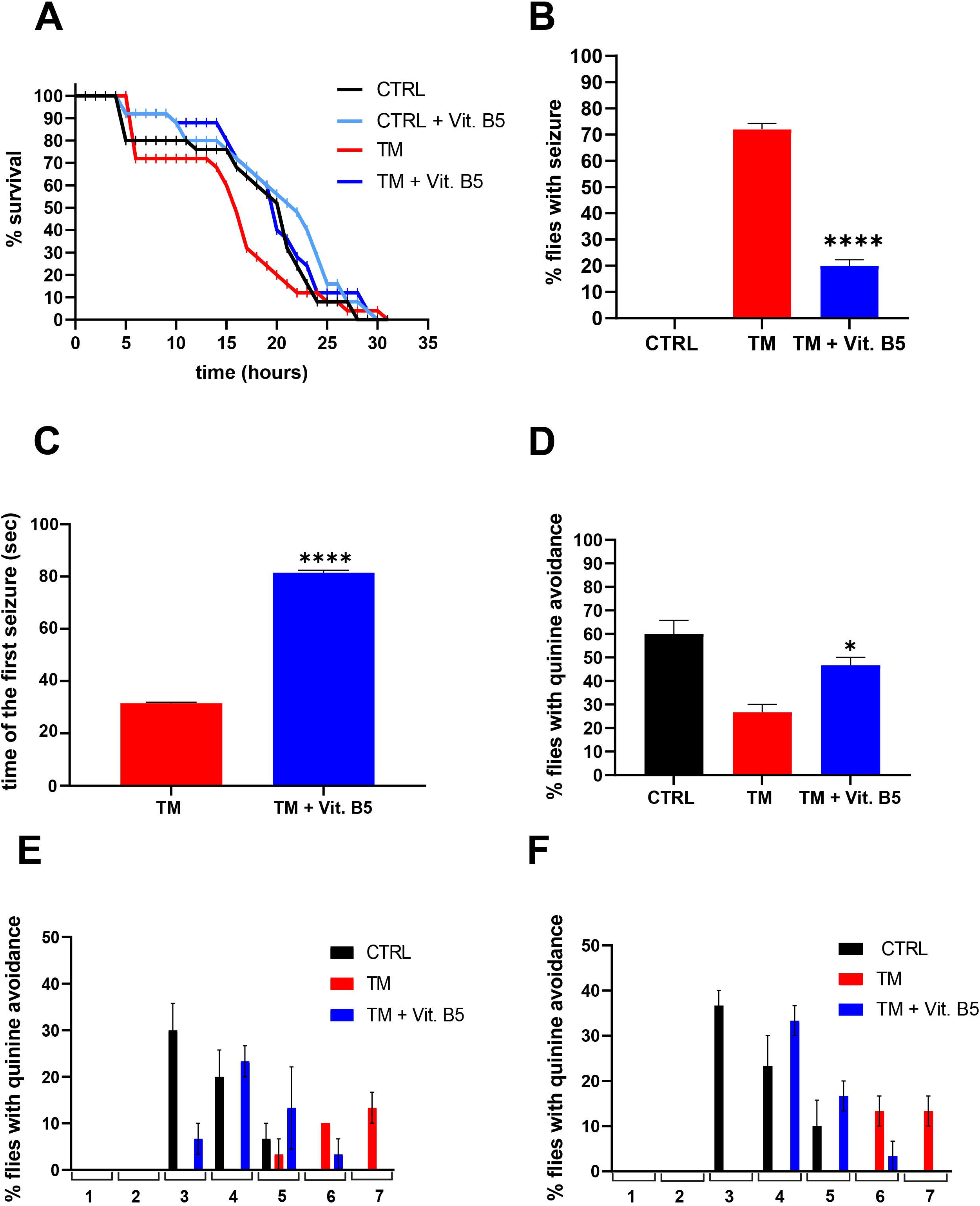
*In vivo* assays with male fly populations. (A) Male fly populations were placed into empty plastic vials and survival was examined every hour until all flies were dead. (B), (C) Male fly populations were subjected to elevated temperatures for 120 seconds as stated in the Methods section. The percentage of flies experiencing a seizure (B) and the time to the first seizure (C) are shown. (D) Male fly populations were subjected to a learning assay as described in the Methods section. The percentage of flies that learned to avoid the quinine-laced light side of the setup were recorded. Vitamin B5 was included in the food as indicated for at least one generation prior to the assays. The concentrations of vitamin B5 used were 4 mM in (A) and 2 mM in (B), (C) and (D). N values for each experiment shown were 25 in (A), 75 in (B) and (C), and 30 in (D). The asterisks for statistical analyses indicate p≤0.05 (*), p≤0.01 (**) and p≤0.0001 (****). CTRL indicates *Ore*^*R*^ flies and TM indicates *TANGO2*^*G517*^ flies. The overall results of the learning trials (see Figure 1D for females and S1D for males) are shown for males (E) and females (F) at each training (1-6) or test (7) cycles. Both male and female *Ore*^*R*^ wild type (CTRL) flies learned avoidance predominantly during the third and fourth training cycles and some in the fifth. In contrast, the *TANGO2*^*G517*^ female flies learned this avoidance during the sixth training cycle and during the testing (cycle 7). When the *TANGO2*^*G517*^ mutants were grown in the presence of vitamin B5, their ability to learn avoidance was observed earlier and predominantly during the fourth, fifth and sixth training cycles.

**Figure S2.**
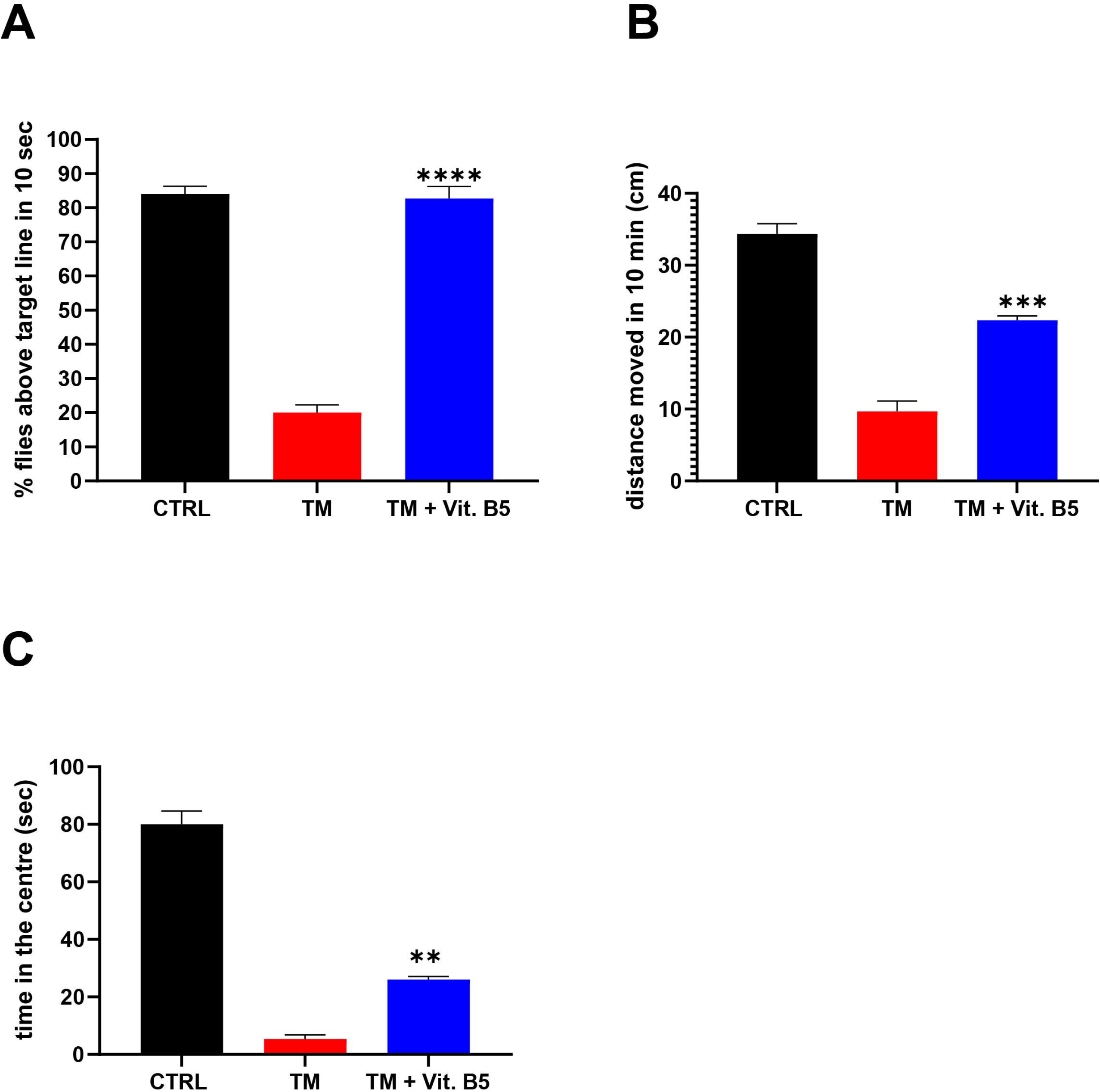
*In vivo* assays with male fly populations. (A) Male flies were placed into an empty vial, gently knocked to the bottom of the vial and were examined as they climbed. The percentages of flies that reached a line 5 cm from the bottom of the vial within 10 seconds are shown. (B) Adult male flies were placed in the center of a 10 cm diameter petri dish and the distance moved over 10 minutes is displayed, as well as the time spent in the central area of the dish (C). Vitamin B5 supplement was included in the food for at least one generation prior to the assays. The concentrations of vitamin B5 used were 4 mM. N values for each experiment shown were 50-75 in (A) and 15-35 in (B) and (C). The asterisks for statistical analyses indicate p≤0.01 (**) and p≤0.0001 (****). CTRL represents *Ore*^*R*^ flies and TM represents *TANGO2*^*G517*^ flies.

**Figure S3.**
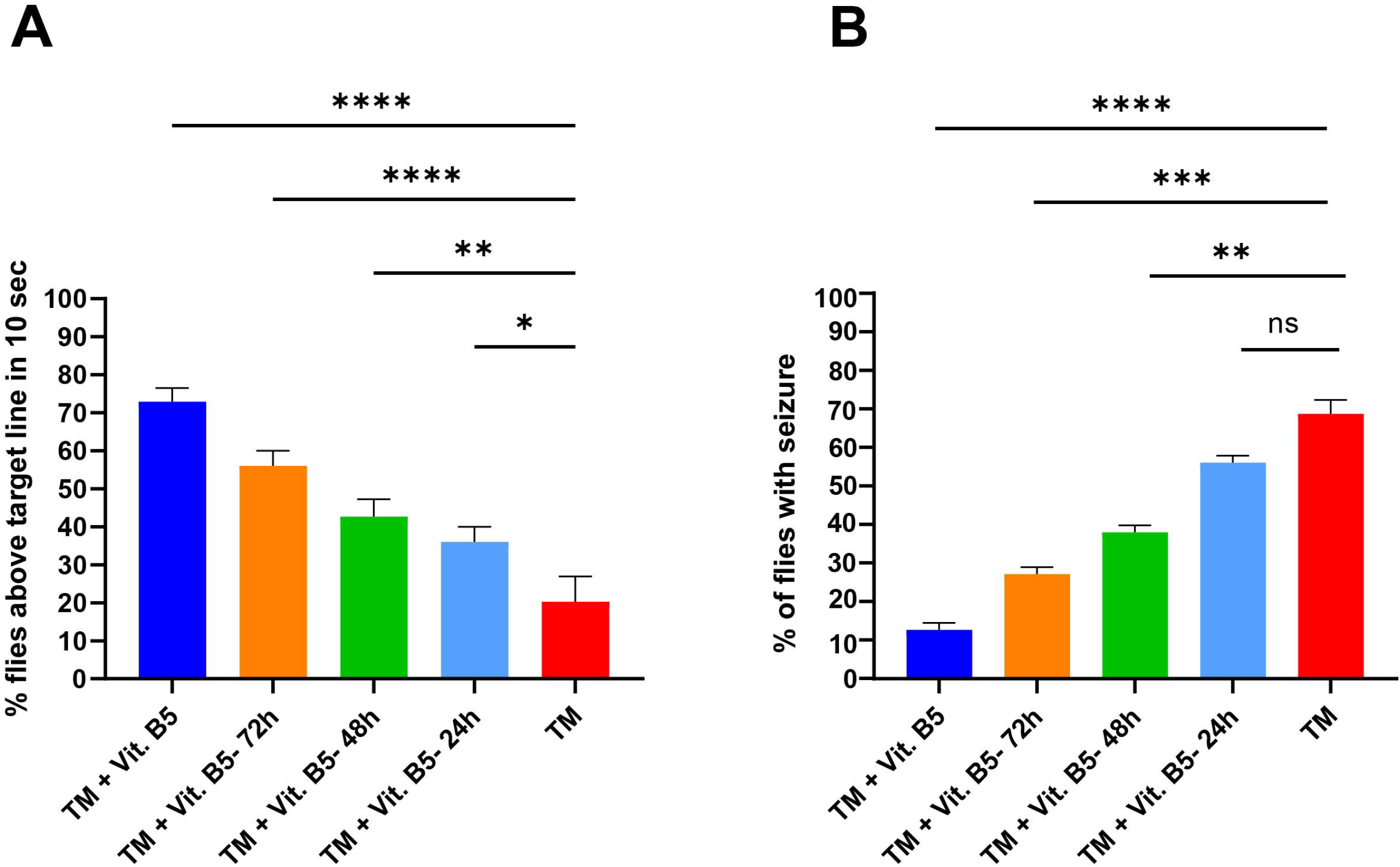
*In vivo* assays with male fly populations. *TANGO2*^*G517*^ male flies were grown either on food without vitamin B5, or grown for at least one generation on vitamin B5-supplemented food. A portion of the untreated adult *TANGO2*^*G517*^ flies were placed on vitamin B5-containing food either 1, 2 or 3 days prior to the assays. Flies were then assayed for their climbing ability (A) and resistance to heat-induced seizures (B). The concentration of vitamin B5 used was 4 mM. N values for each experiment shown were 75 in (A) and 75 in (B). The asterisks for statistical analyses indicate p≤0.05 (*), p≤0.01 (**), p≤0.001 (***) and p≤0.0001 (****). TM represents *TANGO2*^*G517*^ flies.

